# Life stage- and sex-specific sensitivity to nutritional stress in a holometabolous insect

**DOI:** 10.1101/2024.09.02.610820

**Authors:** Leon Brueggemann, Pragya Singh, Caroline Müller

**Affiliations:** Department of Chemical Ecology, Bielefeld University, Universitätsstr. 25, 33615 Bielefeld, Germany; Joint Institute for Individualisation in a Changing Environment (JICE), University of Münster and Bielefeld University, Bielefeld, Germany

**Keywords:** behaviour, energy metabolism, life-history, niche conformance, phenotypic plasticity, sensitive phases, starvation

## Abstract

1. Over the course of their lives, organisms can be repeatedly exposed to stress, which shapes their phenotype. At certain life stages, known as sensitive phases, individuals might be more receptive to such stress than at others. One of these stresses is nutritional stress, such as food limitation. However, little is known about how plastic responses differ between individuals experiencing nutritional stress early versus later in life or repeatedly, particularly in species with distinct ontogenetic niches. Moreover, there may be sex-specific differences due to distinct physiology.
2. The turnip sawfly, *Athalia rosa*e (Hymenoptera: Tenthredinidae), is a holometabolous herbivore, whose larvae consume leaves and flowers, while the adults take up nectar. We examined the effects of starvation experienced at different life stages on life-history traits as well as adult behavioural and metabolic traits to determine which life-stage may be more sensitive to nutritional stress and how specific these traits respond. We exposed individuals to four distinct nutritional regimes, no, larval, or adult starvation, or starvation periods during both larval and adult stage.
3. Larvae exposed to starvation had a prolonged developmental time, and starved females reached a lower initial adult body mass than non-starved individuals. However, males did not differ in initial adult body mass regardless of larval starvation, suggesting the ability to conform well to poor nutritional conditions, possibly through changes in development and metabolism.
4. Adult behaviour, measured as activity, was not significantly impacted by larval or adult starvation in either sex. Individuals starved as larvae had similar carbohydrate and lipid (i.e. fatty acid) contents as non-starved individuals, potentially due to building up energy reserves during their prolonged development, while starvation during adulthood or at both stages led to reduced energy reserves in males.
5. This study indicates that the sensitivity of a life stage to nutritional stress depends on the specific trait under consideration. Life-history traits were mainly affected by larval nutritional stress, while activity appeared to be more robust and metabolism mostly impacted by the adult nutritional conditions. Individuals differed in their ability to conform to the given environment, with the responses being life stage- and sex-specific.

## 1 INTRODUCTION

Sensitive phases are periods during ontogeny when environmental cues, such as temperature, humidity and resource availability, profoundly influence development and subsequent life-history strategies (Fawcett & Frankenhuis, 2015; Walasek et al., 2024). These phases critically shape an individuaĺs morphology, behaviour and physiology (Macdonald, 1985; Monaghan, 2008). Sensitive phases require plasticity and are documented across diverse taxa, including birds, rodents and humans (Oyama, 1979; Baptista & Petrinovich, 1984; Sachser et al., 2020). In organisms that undergo distinct life stages, such as holometabolous insects, sensitive phases may be particularly important, because individual responses to stressors and their general resource requirements are likely to differ (English & Barreaux, 2020). For example, in many insects larvae feed on leaves, while adults feed on nectar; thus, distinct stages show ontogenetic niche shifts (Dopman et al., 2002).

Investigating sensitive phases in holometabolous insects may help us understand population dynamics and evolutionary trajectories under environmental change scenarios and help in developing pest management and conservation strategies (Frankenhuis et al., 2019; Smith et al., 2020).

In nature, individuals may face different stresses throughout their lifespans, with potential for additive effects (Todgham & Stillman, 2013). In response to stressful environments, individuals can alter their phenotype, such as life-history traits, behaviour and the metabolism, leading to niche conformance (Müller et al., 2020). Moreover, individuals may respond differently, leading to individualised niches (Trappes et al., 2022; Kaiser et al., 2024). Recurrent stress exposure across life stages is less studied, but the limited evidence suggests significant impacts on an individual’s phenotype (Gilad et al., 2018; Paul et al., 2019). Stress experienced during sensitive phases may cause more pronounced changes in phenotypes, since the receptivity might be increased (Cheng et al., 2021). Among ecological stressors, nutritional stress, caused by inadequate food quality or quantity (Holmes et al., 2020), is a pervasive challenge faced by organisms in natural environments, exerting significant selective pressure on various life-history traits (Frago & Bauce, 2014). Food limitation may invoke complex developmental, behavioural, and physiological adjustments aimed at maximising fitness during development (Rashid et al., 2021), enhancing foraging efficiency (Scharf, 2016) and prioritising essential physiological processes, such as increased lipid accumulation (Ziegler, 1991; Wang et al., 2016; Yamada et al., 2018). Furthermore, a balance between lipids as long-term energy reserves and carbohydrates as a more rapid, short-term reserve must be achieved (McCue, 2010).

In addition to the life stage, sexes often differ in their responses to nutritional stress during sensitive phases. In *Tribolium castaneum*, females are overall more starvation-tolerant than males (Gilad et al., 2018). However, in many insect species, early-life food stress has been found to have a more negative impact on females than males (Teder & Kaasik, 2023). Differences in sensitivity between sexes may arise if the timeframes of sensitive phases differ, for example, due to varying development times (Rohde et al., 2015) or sexual size dimorphism (Shingleton & Vea, 2023). Sexual size dimorphism may also correlate with sex-specific plasticity, as found in *Drosophila* (Vea et al., 2023), shaping behaviour and metabolic regulation in female and male phenotypes (Shingleton & Vea, 2023). The intertwined sex-specific effects of stress during sensitive phases in insects are still not well understood.

The turnip sawfly, *Athalia rosae* (Hymenoptera: Tenthredinidae), is a holometabolous species characterised by a complex life cycle. The larvae feed on leaves and at later stages also on flowers of Brassicaceae species (Bandeili & Müller, 2010), including crops such as cabbage and rapeseed, while the adults feed on nectar from Apiaceae species. Depletion of nutritional resources can occur at different life stages. For example, high numbers of eggs laid on a plant can cause rapid food exhaustion by the hatching larvae (Saringer, 1976).

Adults may experience nutritional stress due to ephemeral food sources, for example, if they pupated in habitats with suitable food plants that are no longer available at the time of adult emergence (Oishi et al., 1993). Starvation during the larval stage has been shown to prolong the development time and decrease adult body mass but had no effects on adult lifespan of *A. rosae* (Paul et al., 2019; 2022). Starved larvae showed increased activity levels, while starved adults showed reduced activity levels compared to non-starved individuals (Singh et al., 2023). However little is known about the role of larval versus adult starvation with regard to life stage-specific sensitivity and trait-specific responses.

In this study, we investigated the effects of single or repeated nutritional stress events (i.e. starvation) experienced during either the larval and/or adult stage, on life-history traits, behaviour and metabolic traits of adults, to assess the sensitivity of these life stages and identify which traits are most impacted by such stress. We expected the most pronounced effects on individuals that were starved during both larval and adult stage, but intermediate effects for individuals only starved as larvae or adults. We predicted that larval starvation would prolong development time, reduce body mass, and decrease lifespan. With regard to behaviour, we expected lower activity levels in starved adults. With regard to metabolism, we expected the highest levels of lipids in non-starved individuals, as they should have been continuously able to build up energy reserves. Repeatedly starved individuals may rely more on carbohydrates as they cannot build up lipids to the same extend, therefore their lipid contents might be lower. Starvation only during the larval stage may trigger conforming mechanisms, such as an enhanced accumulation of lipids, potentially allowing recovery. We expected stronger effects of early-life food stress on females than on males, consistent with findings on other insect species (Teder & Kaasik, 2023). Overall, we aimed to unravel the complex effects of nutritional stress experienced across ontogeny on the outcome of individualised niches.

## 2 MATERIAL AND METHODS

### 2.1 Rearing of insects and plants

Adults of *A. rosae* were collected during late summer in the surroundings of Bielefeld, Germany. For a continuous rearing, adults were offered potted plants of white mustard (*Sinapis alba*, Brassicaceae) for oviposition. After larval hatching, potted, non-flowering plants of cabbage (*Brassica rapa* var. *pekinensis*, Brassicaceae) were provided as food source. Both plant species were grown from seeds (KWS Saat; Kiepenkerl, Germany) in a controlled environment (greenhouse or climate chamber, about 20 °C, 16L∶8D, 70% relative humidity). Insects were kept in cages (60 x 60 x 60 cm) at a 16:8 h day:night cycle. For pupation, larvae of the last instar, called eonymphs, went into the soil provided in Petri dishes or in the potting soil. Emerging F1 adult individuals were allowed to mate and provided with 2% honey water. Thirty of these females were set up in a new cage with white mustard for oviposition and removed after 48 h from the cage. After five days, 120 freshly hatched larvae (F2) were carefully transferred to individual Petri dishes (5.5 cm diameter) lined with moist filter paper, placed in a climate cabinet under constant conditions (20 °C, L16:D8, 65% r.h.) and used for the experiments. The larvae were offered discs (22 mm diameter) of middle-aged cabbage leaves *ad libitum*.

### 2.2 Starvation treatments and measurements of life-history traits

To study the effects of starvation experience during the larval and/or adult phase on life-history traits, behaviour and metabolism of *A. rosae,* the following starvation treatments were set up (Fig. 1). Half of the larvae were randomly assigned to a larval starvation exposure. From those larvae, cabbage leaf discs were removed twice for 24 h each, once in the third instar and once after having moulted to the fourth instar, but water was still provided through a moist filter paper. All larvae were checked daily for their status (instar and alive/dead). Once individuals reached the eonymph stage, the date was noted and the eonymphs were transferred in individual plastic cups with a gauze lid, containing around 30 g of soil for pupation. Upon adult eclosion, the date was again noted to calculate the developmental time, the sex determined, the individual weighed (initial adult body mass; Sartorius LA 120 S, Germany) and adults transferred into individual Petri dishes. Half of the individuals that had been provided with food *ad libitum* and half of those that had starved as larvae were assigned randomly to an adult starvation exposure, aiming for a balanced sex ratio in each treatment. This resulted in four distinct treatments, individuals that were continuously fed as larvae and adults (NS), individuals that only experienced larval starvation (LS), individuals that only experienced adult starvation (AS) and individuals that experienced repeated starvation during both larval and adult stage (DS). Individuals assigned to an adult starvation exposure were only provided with water on tissue paper for four days. The other adults were provided with a mixture of honey and water (1:10) on tissue paper. After four days of treatment, each individual was weighed again, and the behaviour of a subset of individuals was tracked within the available time constraints (n = 20-25 per treatment group; see Behavioural trials). Following the behavioural trials, half of the individuals of each treatment group (n = 12-15 per treatment group) were frozen at −80°C for analysis of the energy metabolism (see Analysis of energy metabolism). Individuals in the other half of each treatment group were all provided with the honey water mix *ad libitum* and checked daily for survivorship to determine their lifespan.

**FIGURE 1.**
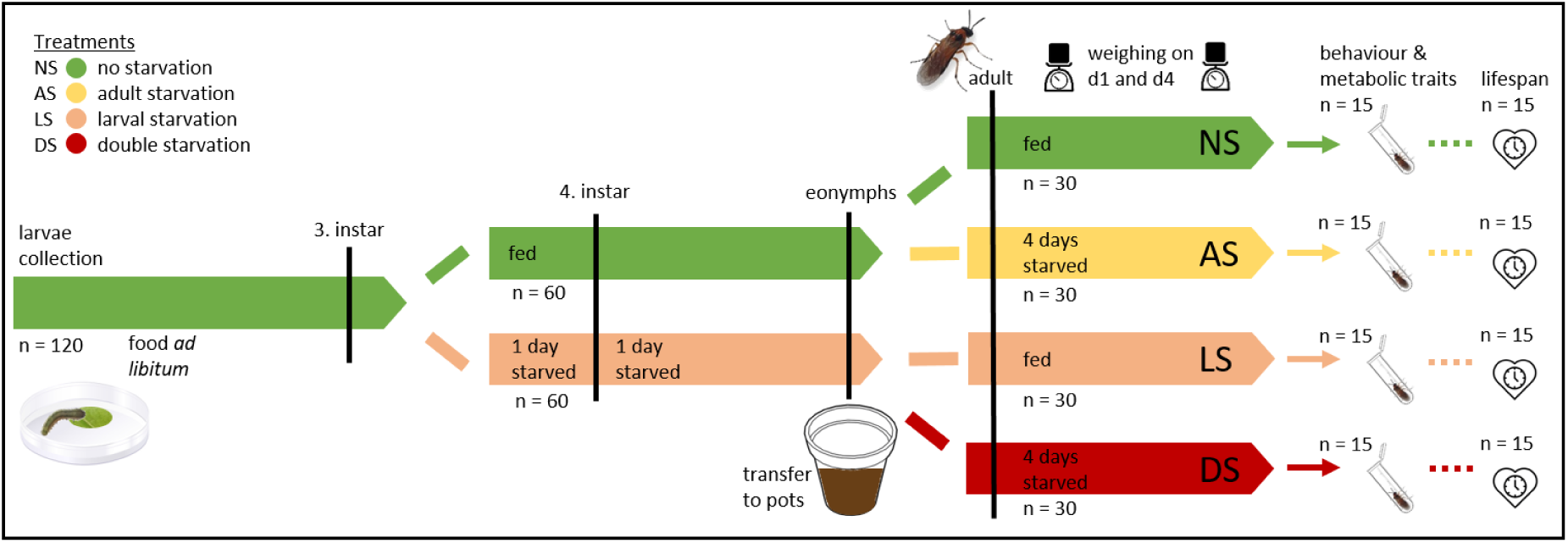
Experimental design for testing sensitive phases to nutritional stress. Individuals of *Athalia rosae* were assigned to different starvation treatments. Data on life-history (development time, body mass, lifespan) as well as behaviour and metabolic traits of adults were collected.

### 2.3 Behavioural trials

Behavioural observations were conducted between 12 pm – 5 pm at 20 °C room temperature. To measure activity levels, adults were transferred with minimal handling to new empty Petri dishes and six individuals were filmed in parallel for 1 h with an overhead camera (Computar, USA). Their activity was tracked using Ethovision V7 (centre-point tracked at 1.92 samples/s; Noldus, Netherlands), extracting the distance moved and the immobility duration (time spent immobile) for each individual.

### 2.4 Analysis of energy metabolism

As measures of energy metabolism, individual levels of lipids (i.e. unsaturated fatty acids) and carbohydrates were analysed using a modified method by Cuff et al. 2021 (Cuff et al., 2021). Frozen sawflies were lyophilised and the dry mass measured. Lipids were extracted by soaking each sawfly in 0.5 ml of 1:12 chloroform:methanol (chloroform: HPLC grade, AppliChem; methanol: LC-MS grade, Fisher Scientific) for 24 h. Afterwards, 250 μl of the supernatant was taken for later lipid determination. As much of the remaining supernatant as possible was discarded and any residual was allowed to evaporate. Then the samples were soaked in the same solvent for another 24 h, but this time all supernatant was discarded, to ensure any residual lipids were removed from the sample prior to carbohydrate extraction. For the carbohydrate extraction, the sawfly samples were homogenised in a mill at 30 Hz for 30 seconds, 0.5 ml of 0.1 M NaOH (VWR International) was added to each sample, samples were incubated on a shaker at 80°C and 250 rpm for 30 min, and then removed and left at room temperature overnight. On the next day samples were centrifuged for 10 min at 13000 rpm and 450 μl of supernatant used for carbohydrate determination.

For the determination of fatty acids, which make up about 50 % of the lipids in Hymenoptera (Aguilar 2021), the respective samples (50 μl) were each mixed with 10 μl of sulphuric acid (95%, VWR International) in a well and incubated at 100°C for 10 min. After a 5 min cool down, 240 μl vanillin in phosphoric acid (17% in ultrapure water, AppliChem) was added to each well and the absorbance measured at 490 nm in a microplate reader (Multiskan FC, Thermo Scientific, USA) (adapted from (Cheng, Zheng & VanderGheynst 2011). As a standard dilution series for calibration, a cricket oil (Acheta Cricket Oil, Thailand Unique) in 1:12 chloroform:methanol was measured on the same 96 well plate in eight concentrations.

For carbohydrate determination, sugars were measured, calibrated by trehalose and glycogen, as these are the most abundant sugars in insects (Jiang et al., 2019). The respective samples (40 μl) were each mixed with 160 μl of anthrone reagent (2 mg/ml anthrone (Roth) in 95% sulphuric acid), incubated for 10 min at 92°C, and measured at 620 nm (adapted from (Dreywood 1946). For carbohydrate calibration, a standard dilution series of 1:1 trehalose:glycogen (trehalose: 98%, Roth; glycogen: AppliChem) in ultrapure water was used. Each 96-well plate had four solvent blanks and two calibration rows.

### 2.5 Statistical analysis

Statistical analyses were conducted using the statistical software R, Version 4.1.3 (R Development Core Team 2010). For the replication statement, refer to table 1. To compare the larval development time and initial adult body mass between the individuals non-starved and starved as larvae, a Mann Whitney U-test was used. For comparing adult traits (i.e. body mass change, distance moved, time spent immobile, lipid and carbohydrate mass) among the four treatment groups, a Kruskal-Wallis test was used, followed by post-hoc pairwise comparisons using Dunn’s test with Bonferroni correction. Distance moved and time spent immobile were tested for correlation with a Spearman rank correlation. Survival probability of individuals was plotted with Kaplan-Meier curves using the survival package (Therneau 2024) and an overall log-rank test was performed, followed by pairwise log-rank tests with Bonferroni-Holm correction of *p*-values. Graphical visualisations were generated using the ggplot2 (Wickham 2016) multcompView, ggthemes and ggdist packages.

**TABLE 1.** Replication statement. Overview of scales and sample sizes for recorded traits. NS = no starvation, LS = larval starvation, AS = adult starvation, DS = starvation as larvae and adults.

| Scale of interference | Scale at which the factor of interest is applied | Number of replicates at the appropriated scale |
| --- | --- | --- |
| Insect (*A. rosae*) | Individuals for life-history traits | NS= 13 ♀, 16 ♂; LS = 10 ♀, 18 ♂;<br>AS = 13 ♀, 17 ♂; DS = 10 ♀, 19 ♂ |
| Insect (*A. rosae*) | Individuals for behaviour | NS = 7 ♀, 13 ♂; LS = 10 ♀, 18 ♂;<br>AS = 5 ♀, 16 ♂; DS = 10 ♀, 18 ♂ |
| Insect (*A. rosae*) | Individuals for metabolism | NS = 5 ♀, 10 ♂; LS = 6 ♀, 9 ♂;<br>AS = 4 ♀, 11 ♂; DS = 4 ♀, 10 ♂ |

## 3 RESULTS

### 3.1 Larval starvation prolonged larval development and reduced female initial adult mass

Starvation during the larval stage led to a significantly prolonged development for both sexes (females *U* = 24, males *U* = 66; *p* < 0.001), with starved larvae taking nearly two days longer than non-starved ones (Fig. 2A). Two females and one male individual from the larval starvation treatment took one additional instar to reach the eonymph stage. The initial adult body mass differed significantly for females (*U* = 366, *p* = 0.019) but not males (*U* = 662, *p* = 0.549), with females starved as larvae being significantly lighter than non-starved females (Fig. 2B).

**FIGURE 2.**
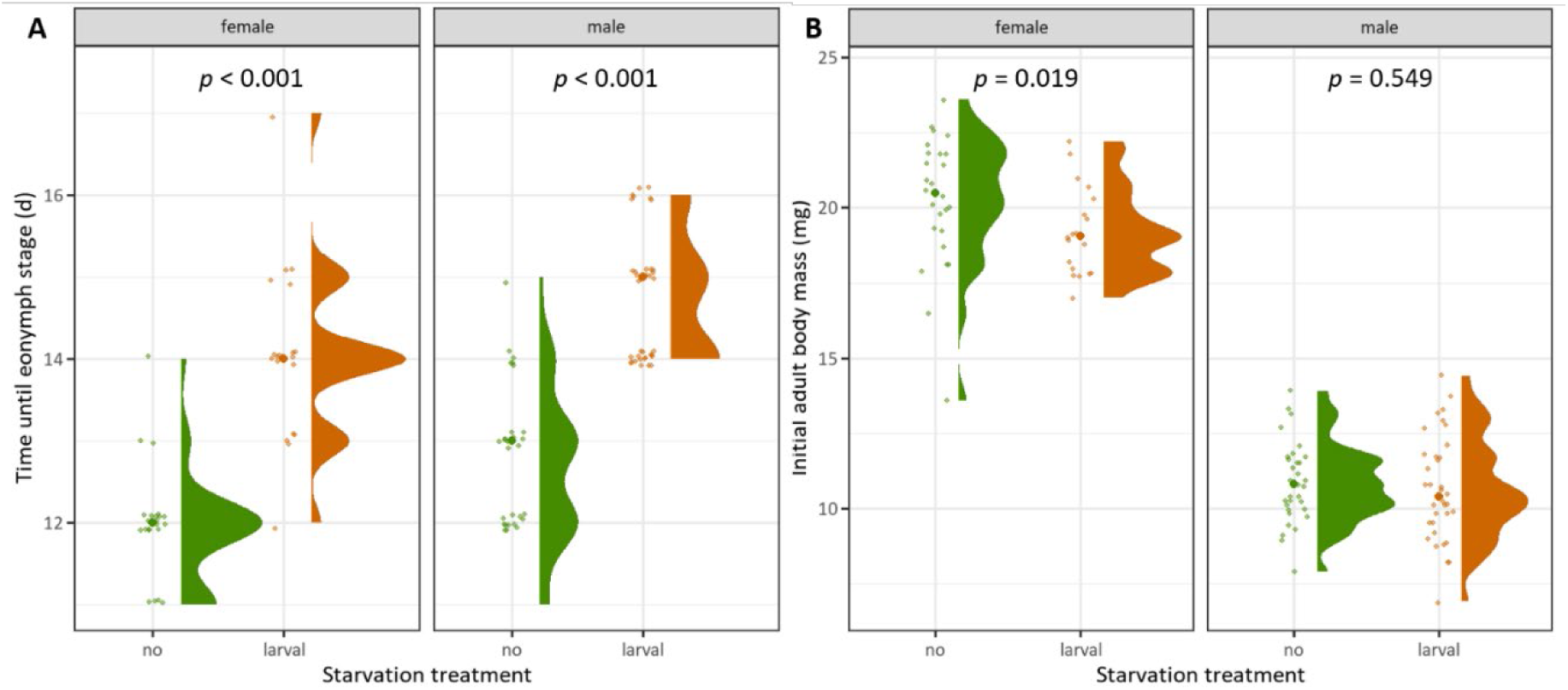
Effect of larval starvation on A) larval development time until eonymph and B) initial adult body mass (fresh weight). Half violin plots indicate the distribution of values, raw data is plotted with slight jittering and thicker dots indicate medians. Statistical differences were tested with Mann-Whitney U-tests.

### 3.2 Adult and repeated starvation reduced female body mass with no effect of starvation treatments on lifespan

Treatment had a significant effect on adult body mass change in females only (females: *ꭓ^2^*= 24.11, *df* = 3, *p* < 0.001; males: *ꭓ^2^* = 4.826, *df* = 3, *p* = 0.186). Females starved as adults or as both larvae and adults had a significantly lower body mass change compared to those in the other starvation treatments (Fig. 3).

**FIGURE 3.**
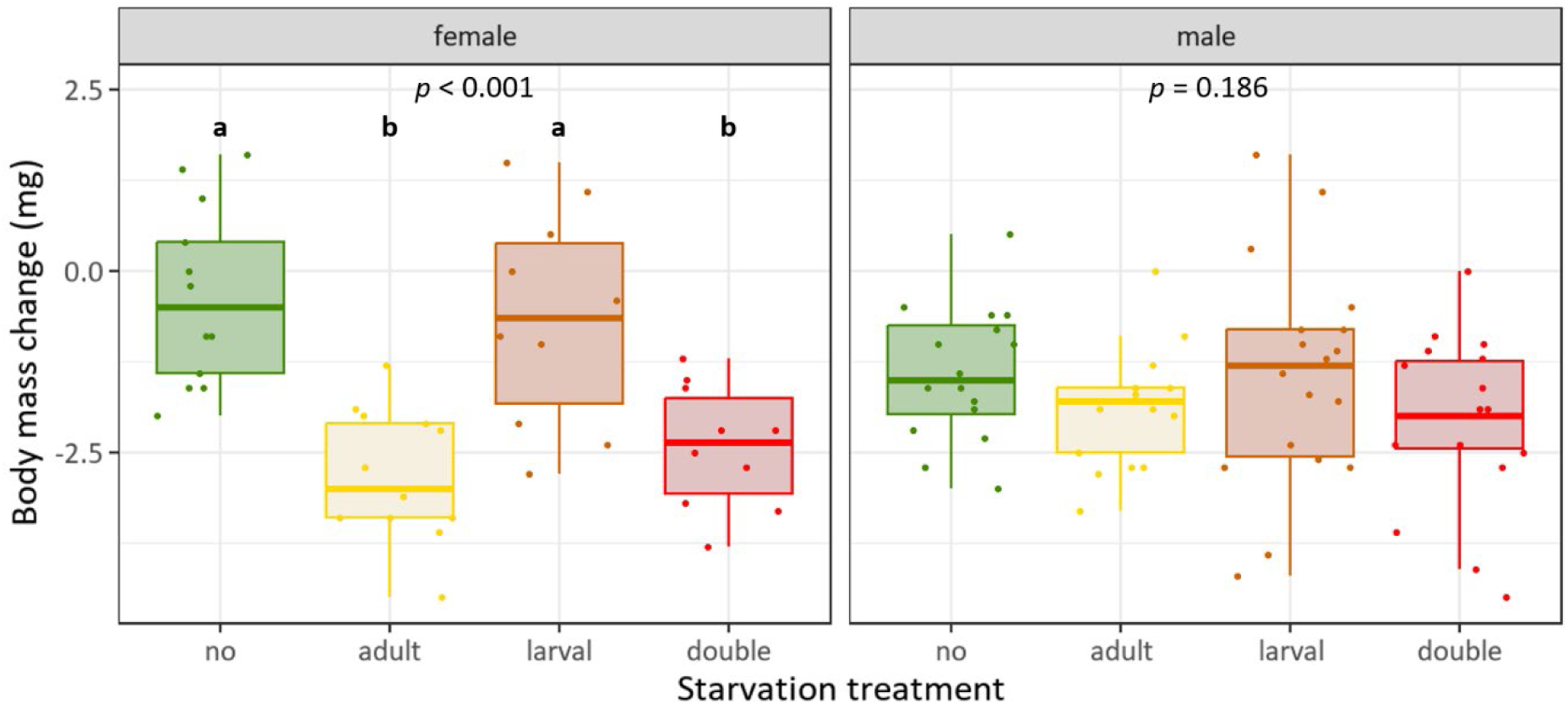
Effects of larval and/or adult starvation on body mass change (fresh weight) within 4 days in female and male adults. Boxplots indicate interquartile range (IQR, boxes) with medians, whiskers extent to ±IQR*1.5, dots are raw data points. Overall significant differences were tested with Kruskal-Wallis rank sum test, followed by pairwise Dunn posthoc test with Bonferroni correction. Different letters denote significantly different (*p* ≤ 0.05) treatment effects.

Adult survival probability, i.e. lifespan, was not affected by treatment for both females (*ꭓ^2^* = 1.4, *df* = 3, *p* = 0.7) and males (*ꭓ^2^* = 7.2, *df* = 3, *p* = 0.066). However, males starved at both life-stages showed a trend toward lower survival probability (Fig. 4).

**FIGURE 4.**
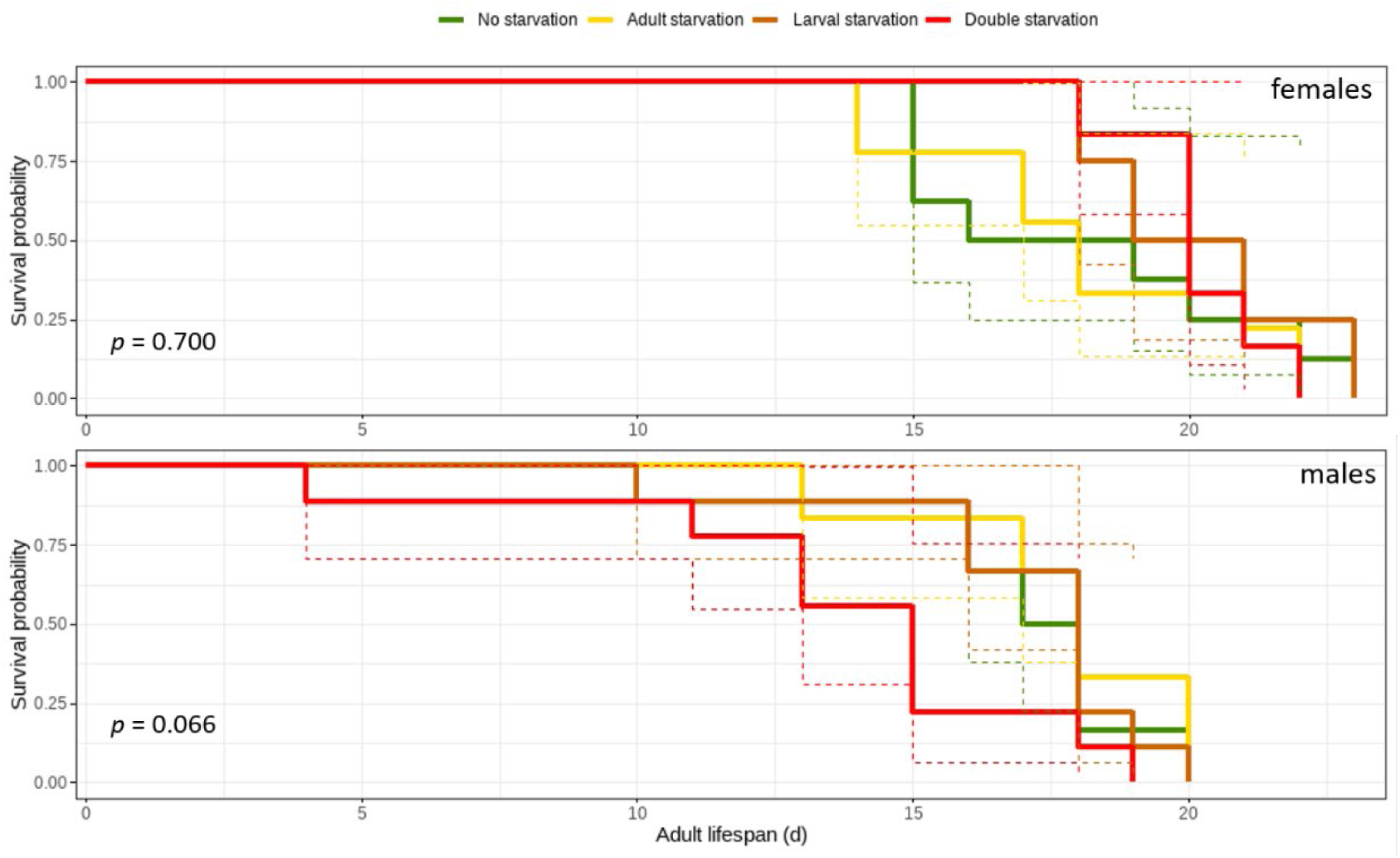
Effect of larval and/or adult starvation on female and male adult survival probability, represented by Kaplan-Meier curves. Dashed lines show 95% confidence intervals. An overall log-rank test was performed, followed by pairwise log-rank tests with Bonferroni-Holm correction of *p*-values.

### 3.3 No effect of starvation treatments on behaviour

There was no significant effect of treatment on either distance moved (females: *ꭓ^2^* = 4.416, *df* = 3, *p* = 0.220; males: *ꭓ^2^* = 0.547, *df* = 3, *p* = 0.908; Fig. 5A) or immobility duration (females: *ꭓ^2^* = 5.702, *df* = 3, *p* = 0.127; males: *ꭓ^2^* = 0.273, *df* = 3, *p* = 0.965; Fig. 5B) in either sex. The two traits were significantly correlated across all treatments (all *r* ≤-0.931 and *p* <0.001).

**FIGURE 5.**
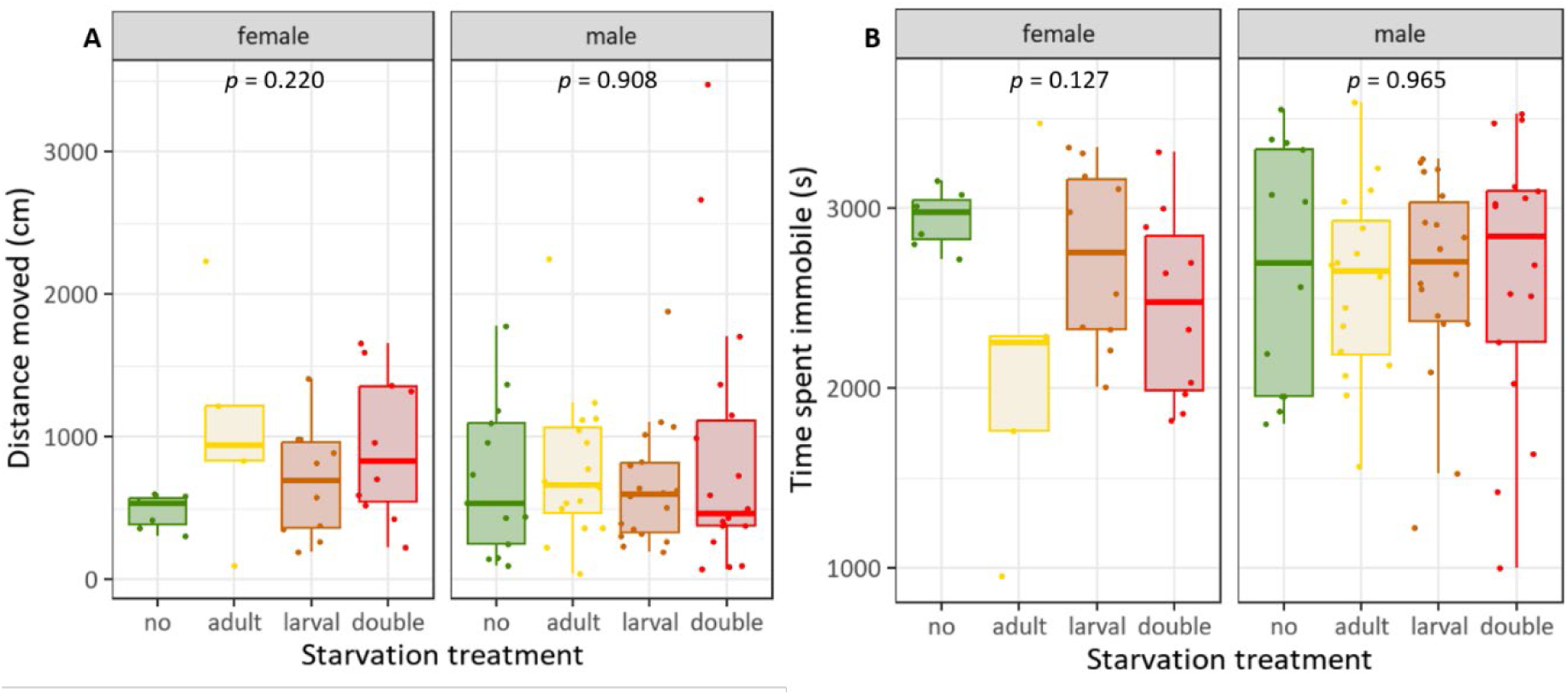
Effects of larval and/or adult starvation treatments on A) distance moved and B) time spent immobile during 1hour of recording of adults. Boxplots indicate interquartile range (IQR, boxes) with medians, whiskers extent to ±IQR*1.5, dots are raw data points. Overall significant differences were tested with Kruskal-Wallis rank sum test, followed by pairwise Dunn posthoc test with Bonferroni correction.

### 3.4 Adult and repeated starvation led to a decrease in both lipids and carbohydrates in males

The lipid mass per body mass was significantly impacted by treatment in males only (females: *ꭓ^2^* = 3.122, *df* = 3, *p* = 0.373; males: *ꭓ^2^* = 12.793, *df* = 3, *p* = 0.005), with individuals having starved in both the larval and adult stage showing lower lipid amounts compared to non-starved individuals and to individuals starved only during the larval stage (Fig. 6A). The carbohydrate mass per body mass was significantly affected by the starvation treatments in both sexes (females: *ꭓ^2^* = 7.908, *df* = 3, *p* = 0.048; males: *ꭓ^2^* = 9.137, *df* = 3, *p* = 0.028), although post hoc comparisons showed no significant differences between treatments for females (Fig. 6B). In males, those starved as adults or both as larvae and adults had lower carbohydrate levels compared to non-starved individuals or those starved only during the larval stage.

**FIGURE 6:**
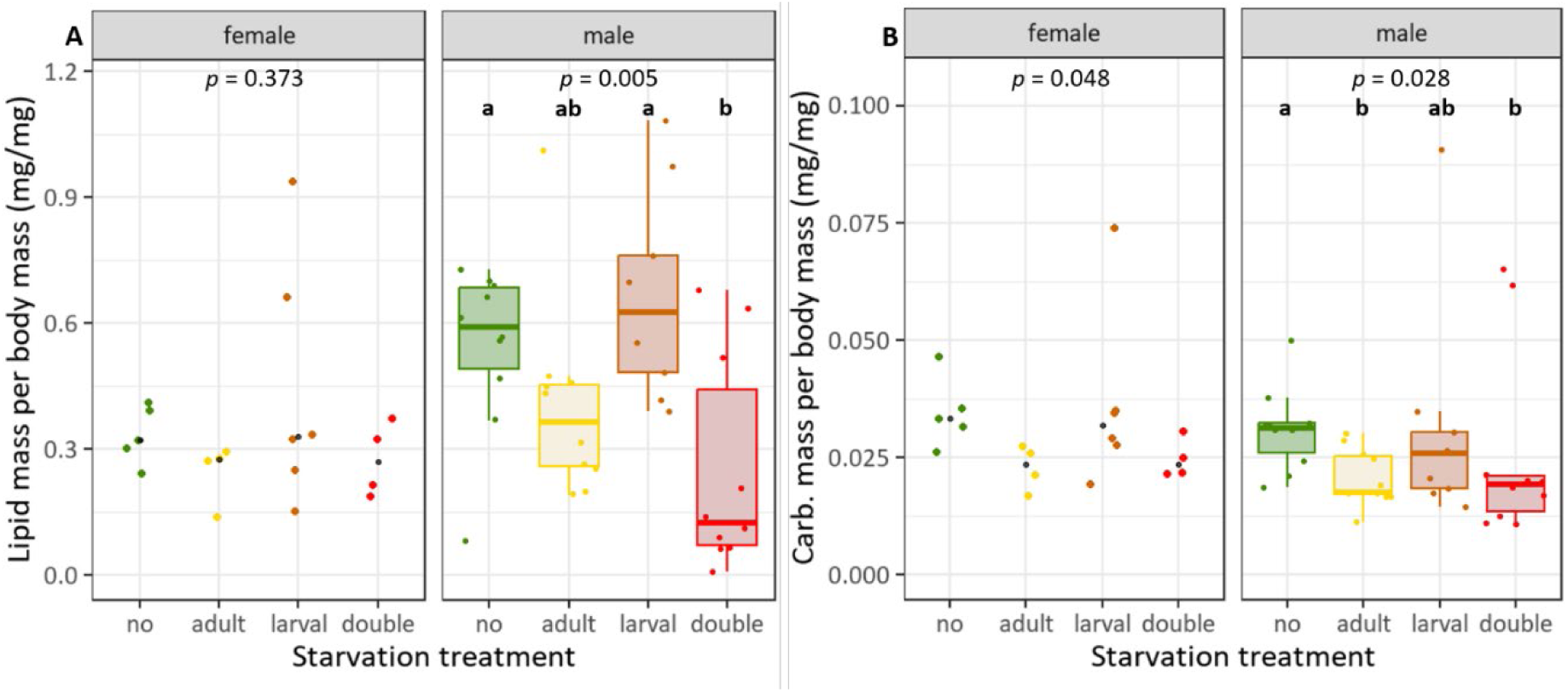
Effects of larval and/or adult starvation treatments on energy metabolism with A) lipid content per body mass and B) carbohydrate content per body mass (dry weight) of adult sawflies. Boxplots indicate interquartile range (IQR, boxes) with medians, whiskers extent to ±IQR*1.5, dots are raw data points. Overall significant differences were tested with Kruskal-Wallis rank sum test, followed by pairwise Dunn posthoc test with Bonferroni correction. Different letters denote significantly different (p < 0.05) treatments. Black dots indicate the median for females. Boxplots are shown only for males due to n = 4 in two female treatment groups.

## 4 DISCUSSION

By recording a comprehensive set of life-history, behavioural and metabolic traits, we aimed to identify phases during which individuals are most sensitive to starvation. Our findings show that these phases may differ depending on the respective trait.

Starvation during the larval stage led to prolonged development until reaching the eonymph stage and also sometimes additional larval instars. Similar effects were also found in response to different starvation regimes, partly with more frequent starvation events during the larval stage, in the same species (Paul et al., 2019; 2022). With an additional instar, individuals may compensate for inadequate nutrition and reach a larger adult body mass at the expense of a prolonged larval development (Esperk et al., 2007). Delayed development may result from the lack of needed nutrients and is probably also connected to delayed growth hormone production (Friend, 1958; Clark & Gibbs, 2023). It is well-documented across many insect taxa that juvenile/larvae exposed to lower resource levels significantly prolong their developmental time and remain smaller compared to conspecifics in more favourable conditions (Teder et al., 2014). Moreover, consistent with our expectation and findings from other insect species (Teder & Kaasik, 2023), we found that only adult females were lighter at eclosion as a result of larval starvation. In contrast, males seemed to be able to compensate for the poor larval conditions, reaching comparable adult body mass as non-starved larvae. This sex-specific difference in adult body mass in response to larval starvation is also in accordance with previous findings on *A. rosae* (Paul et al., 2022). The development of female reproductive organs might be more demanding and prioritised, restricting nutrient availability for biomass production (Jensen et al., 2015). Males may compensate the effect on this trait well by prioritising essential cells and tissue (Metcalfe & Monaghan, 2001). Contrary to that, other studies found that females of holometabolous insects may be more plastic in their body size (Rohner et al., 2018).

When adults were exposed to starvation, again only females showed a significant loss in adult body mass, with no signs of an additive effect of larval and adult starvation. Rather, the body mass loss was comparable for individuals starved at both stages or only as adults, which may indicate that individuals that experienced starvation already during early life may have adjusted their metabolism to starvation. These individuals may gain improved digestive efficiency, as has been found in *Drosophila* (McCue et al., 2017). Adult lifespan was not affected by any of the starvation treatments, indicating that *A. rosae* is quite robust even to repeated starvation events as implemented in our study. However, males starved as larvae and adults showed the lowest lifespan, though this was not statistically significant.

Individuals may show different insulin-signalling and nutrient allocation, as demonstrated in *Drosophila* (Magwere et al., 2004), and therefore react with different sensitivity in terms of lifespan. In conclusion, certain traits may be more affected and sensitive to starvation during specific life-stages than others. During the larval stage, plasticity may help individuals to adapt their metabolism and cope better with subsequent nutritional stress. In contrast, adult body mass may respond more sensitively to starvation during the adult stage than during the larval stage.

In contrast to our expectation, neither larval nor adult starvation affected the activity of the adults, measured as distance moved and time spent immobile. These two traits were negatively correlated. Different effects of starvation on arthropod behaviour have been reported, ranging, for example, from predation avoidance to increased foraging (Scharf, 2016). The life stage at which starvation occurs and the behavioural assays are performed can be important factors (Romero et al., 2010). During early life starvation, insects may increase their activity in order to locate new food sources. In *Drosophila melanogaster*, such hyperactivity has been found to be induced by octopamine (Yang et al., 2015). At a more severe level of starvation, activity may be reduced to conserve energy or due to exhaustion (Scharf, 2016). We did not test for flight capacity, but inadequate nutrition may reduce flight capacity by causing changes in muscle molecular composition (Portman et al., 2015). Activity-related behaviours seemed to be more conserved and less plastic, possibly to maintain key survival functions. This is in contrast to a previous study, in which significant impacts of starvation on activity of *A. rosae* were found (Singh et al., 2023). Different outcomes may result from slight differences in the set-up, such as keeping individuals in groups (Singh et al., 2023) vs. keeping them individually (present study) during the rearing phase. Nevertheless, we cannot exclude that other behaviours, i.e. those related to boldness, may be more responsive to the starvation regimes.

In our energy metabolism assay the sex-ratio was male-biased, so we mainly discuss the effects on males, although the observed pattern was similar for both sexes. Lipid levels significantly differed between males from the distinct treatments. Males with no starvation and those exposed to larval starvation showed higher amounts of lipids per body mass than those in the other two treatments, as hypothesised. Upon larval starvation, individuals may build up an enhanced lipid storage. In the grasshopper *Schistocerca americana*, larval nutrition has been found to be particularly relevant for adult lipid storage, with lipid stores being more important than carbohydrate or protein stores (Hahn 2005). In *A. rosae*, the carbohydrate levels showed a somewhat similar pattern to the lipids in response to the treatments, indicating that both energy sources may be equally important for the adults. The mobilisation of more than just one energy source is essential to cope with nutritional stress (Sanathoibi & Keshan, 2019). For instance, honey bees starved as larvae improve their metabolic response to adult starvation mainly through carbohydrates (Wang et al., 2016). Different species may thus preferably use different energy sources. Larval starvation alone may lead to niche conformance, with individuals building up energy reserves, whereas adult starvation and repeated starvation in the larval and adult stage may lead to an increased depletion of both energy sources. Sensitive phases in insects, such as the larval stage with higher plasticity, can have lasting impacts on the adult physiology and the ability to manage energy resources.

## 5 CONCLUSION

Our study revealed life-stage and sex-specific effects of nutritional stress on various parameters involving life-history traits, behaviour and metabolism. The effects differed across the recorded traits. Life-history traits were affected most significantly, especially in females. Activity-related behaviour seemed to be a more conserved trait in our study, while energy metabolism was mainly affected by adult starvation and starvation during the larval and adult stage. Our findings indicate that the sensitivity of a certain life stage to nutritional stress depends on the specific trait under consideration, and is not uniform across all traits in a single life stage. The larval stage appears more sensitive with respect to life-history traits, whereas the adult stage seems more sensitive in terms of metabolism. Additionally, the intensity and frequency of stressors are decisive. Being sensitive with regard to specific traits in different stages may lead to distinct individualised niches.

## Acknowledgements

We thank Zacharias Ampofo for helping with general maintenance and behavioural assays, Emma Cyplik for maintaining the *Athalia rosae* laboratory culture, and the gardeners of Bielefeld University and Karin Djendouci for rearing cabbage and mustard plants.

